# A photo-switchable gold nanoformulation based on the dCas9 protein for spatiotemporal controlled gene editing activation in vivo

**DOI:** 10.1101/2024.06.13.598872

**Authors:** Soultana Konstantinidou, Mafalda A. Rocco, Francesco Nocilla, Doriana Debellis, Marta d’Amora, Francesco De Angelis, Chiara Gabellini, Francesco Fuso, Francesco Tantussi, Vittoria Raffa

## Abstract

Genome editing allows for the manipulation of genomic DNA for biotechnology and biomedical applications, but the specificity and control of the editing process remain a challenge. This study introduces a nano-switch for spatiotemporal control of the editing process. The basic module of the nano-switch (the monomer) is composed of a gold nanorod conjugated with the dead Cas9. Based on mathematical models, we established the design and the mechanism of action of the nano-switch. Briefly, when two monomers, guided by their respective guide RNAs, form a dimer onto the DNA and get irradiated with a Near-Infrared pulsed laser resonant at the plasmonic properties of the dimer, they generate a localized heat that triggers a thermal break onto the DNA. The nano-switch was generated, validated, and tested in zebrafish embryos at the 1-cell stage. Molecular analysis of irradiated embryos showed targeted DNA mutations, validating the efficacy of the nano-switch as a tool for conditional gene editing that integrates the if-when-where functions.

## INTRODUCTION

A big revolution in synthetic biology is the ability to rationally engineer intracellular switches that can trigger different signalling cascades in specific cell types within living organisms, given an input signal. The design principles of synthetic biology find important applications in the field of genome editing. This field experienced a major revolution with the discovery of the CRISPR/Cas9 system^1, 2^ This system involves the Cas9 enzyme and a single guide RNA (gRNA), which allows for DNA recognition, targeting, and editing ^3^. Despite its high efficiency and broad applicability, Cas9 can tolerate mismatches between the gRNA and the target sequence in the genome, resulting in off-target effects^4, 5^. In large genomes, the risk of unintended cleavage events by Cas9 at DNA loci different from the target locus is significant, leading to unwanted mutations. To exploit the therapeutic potential of the CRISPR/Cas9 system, there is an urgent need for spatiotemporal control of the editing process to limit or prevent unwanted cuts. This requires the design of a switch that generates the proper response when a number of conditions are met. In other words, the switch would implement multi-input AND Boolean logic gates. In this context, the CRISPR/Cas9 system can be understood as a 1-input logic gate where the output (gene editing) occurs when the input (locus recognition by the gRNA) is true. Adding more input terminals to a logic gate increases the number of input state possibilities (2^n, where n is the number of inputs). AND logic is active if and only if all inputs are on. So, when the number of possible states increases, the probability of a random activation in an unwanted condition is greatly reduced.

Some 2-input AND logic gates have already been proposed. Spatial control can be achieved by using a dimeric nuclease. This can be done by fusing the catalytically dead Cas9 (dCas9) to the FokI nuclease, which works as a dimer, meaning that cleavage requires 2 inputs to occur, i.e., the simultaneous association of two dCas9-FokI monomers to target sites that are 15-25 base pairs apart^6^. Temporal control has been achieved by employing light activation, such as ultraviolet-visible and far-red light, to regulate the temporal reconstruction of Cas9 in the cell^7-10^. In this way, the Cas9 protein is split into portions that reconstitute the functional protein under light.

Here, we propose a nano-switch that implements a 3-input AND logic gate for simultaneous spatiotemporal control of gene editing (Figure 1). The nano-switch exploits the plasmonic behavior of gold (Au) nanomaterials. It is employed as a dimer composed of the dCas9:gRNA linked to an Au nanorod (NR) (Figure 1A). When the two AuNR-dCas9:gRNA monomers bind to target sites at a 9 base pairs distance, the dimer is constituted. When two plasmonic nanomaterials are brought into proximity, the coupling between them increases, and the optical behavior of the dimer prevails if the laser wavelength is centred on the localized surface plasmon resonance (LSPR) of the dimer rather than that of the single monomer. The nano-switch has been designed so that each monomer hybridizes to its cognate target genomic locus and, under the condition of “resonant laser on,” generates a strong confined electromagnetic field in the nanoscopic gap between the two. This strong electromagnetic field also produces rapid and localized heating of the surface of the material and a dramatic temperature jump. A thermally-induced DNA double-strand break would occur with a time constant below milliseconds.

**Figure 1.**
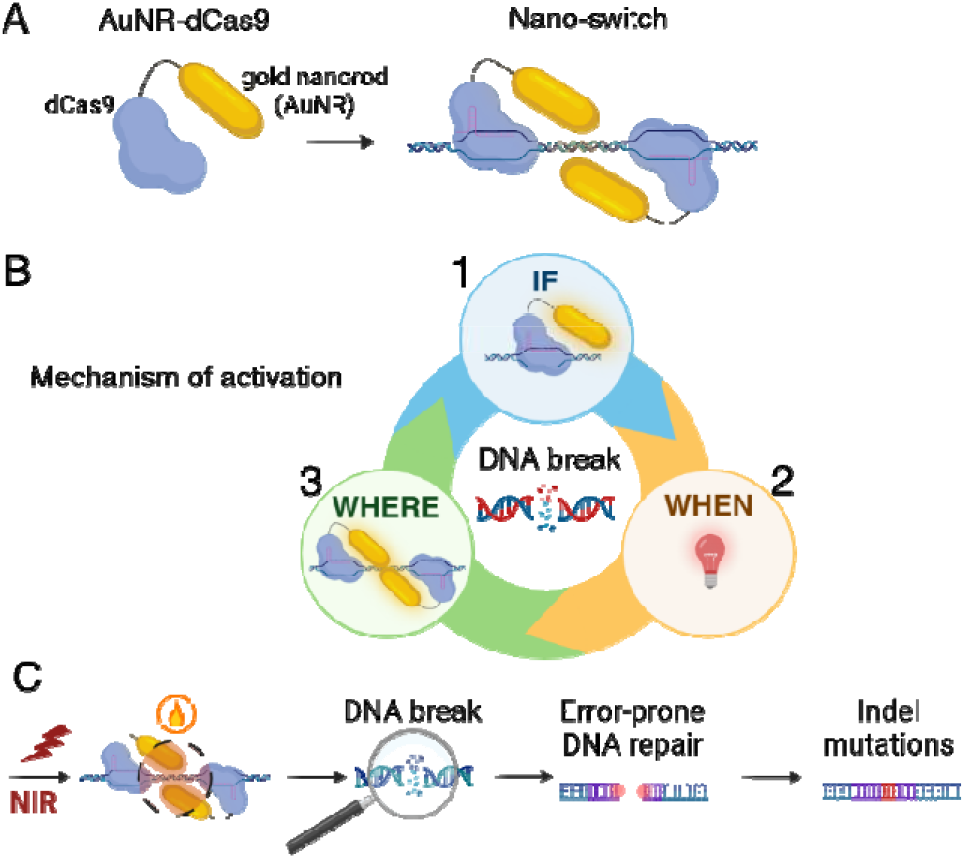
Schematic illustration of (A) the nano-switch, (B) the mechanism of activation of the nano-switch by the 3-input AND logic gate: if-when-where, (C) the editing induced by the thermo-inducible double strand DNA break by the nano-switch activation.

To summarize, the proposed nano-switch implements an if-when-where function, meaning that gene editing occurs if the ribonucleoprotein binds to the locus, when the laser is resonant, and where the dimerization occurs (Figure 1B). We tested the nano-switch in zebrafish at early developmental stages and analyzed for editing modifications occurring from the error-prone repair of the double-strand DNA break (DSB) at 5 hours post-fertilization (Figure 1C).

## RESULTS AND DISCUSSION

The design of the nano-switch was based on the optimization of nanomaterial structure and parameters by numerical analysis. Commercially available COMSOL Multiphysics® software was used to evaluate the optical properties of the AuNRs (absorption, scattering, local electric field intensity, and electromagnetic power loss density) and to estimate their local heating exploited for inducing DSB on DNA. Based on numerical analyses, the optimal conditions to obtain a strong temperature increment with extreme spatial localization correspond to a dimer composed by two gold nanoparticles with a nanorod shape and size of 5×20 nm. The irradiation conditions are ultrashort pulses (200 fs) with a repetition rate of 80 MHz and average power density impinging on the particle in the order of 2·10^7^W/m^2^ (peak power density: D_pp_ = 10^12^ W/m^2^). Figure 2 shows the temperature profile at the surface of the two nanorods (see also video, supporting information) at two different time points (t1, t2 are the time values at the center and end of the 200 fs pulse). According to the simulation, the temperature peak is at the center of each nanorod. The generation of two “hot spots” is believed to be particularly efficient for the generation of a gene knock-out. As the gap between the two monomers decreases, absorption tends to peak at higher wavelength values (Figure 2B). The monomer (represented here as a gap distance of 100 nm) is substantially out of resonance at 870 nm, ensuring optimal suppression of off-target phenomena. According to our simulations, the pair of gRNAs should be designed to have an optimal distance between the nanorods in the range of 5-10 nm. As the nanorods are subjected to Brownian motions, the data in Figure 2 plot the ideal configuration in terms of Poynting vector, polarization of the incident light, and dimer axis. However, even if the.probability that a single 200 fs pulse will be effective is negligible, we can easily assume that with irradiation lasting a few seconds, at least 1 out of 10^8^ pulses will find the nano-switch in the optimal configuration.

**Figure 2.**
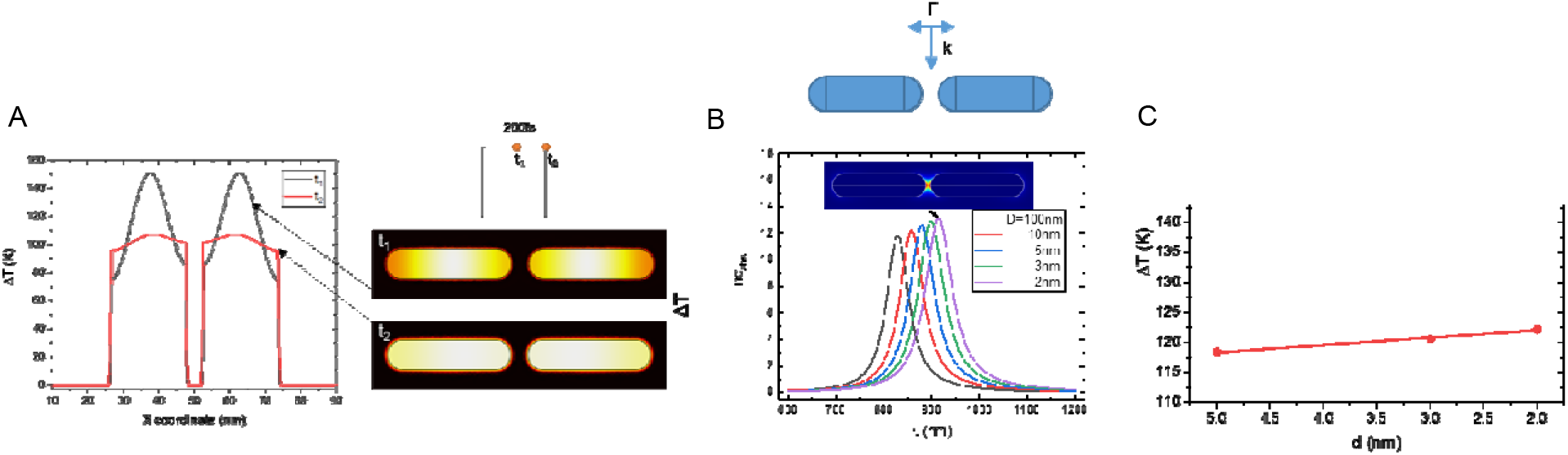
(A) ΔT profile at time instants t1 and t2. 200 fs laser pulse. D_pp_ = 10^12^ W/m^2^@200 fs; 80 MHz repetition rate. (B) Dimer absorption profile as a function of the gap between the monomers. Wave vector, polarisation of the incident light, and the dimer axis used during the simulation are schematically shown. (C) ΔT as a function of the gap between the two monomers (mean value).

As the next step, we synthesized the nano-switch consisting of a gold nanorod (5×20 nm) conjugated with the enzyme dCas9, named AuNR-dCas9. To validate the nano-switch, we needed to demonstrate that the conjugation method does not alter the protein activity. Since dCas9 is a mutated protein without the ability to induce DSB on DNA, we first conjugated the Cas9 protein onto gold nanorods because its endonuclease activity can be easily monitored. The tested conjugation methods were either N-Hydroxysuccinimide (NHS) chemistry or Nitrilotriacetic acid (NTA)-Ni2+-Histidine tag affinity, and the nanoformulation was named AuNR.NHS-Cas9 or AuNR.NTA-Cas9, respectively (Figure 3A). While NHS chemistry potentially targets all amines localized in the lateral chains of protein amino acids, the histidine tag offers the advantage of unidirectional and reproducible binding. The endonuclease activity of the nanoformulations was tested *in vitro* on a linear DNA fragment. Despite the fact that the same concentrations of protein (50, 100, and 150 nM) were tested for both constructs, it was noticed that the AuNR.NTA-Cas9 was more effective than the AuNR.NHS-Cas9 (Figure 3B, C). This may suggest that the orientation of the bond between the Cas protein and the AuNR plays an important role in enabling Cas9 to maintain its catalytic activity after conjugation. Thus, for further experiments, the AuNR.NTA-Cas9 was preferred and named simply AuNR-Cas9. We tested the AuNR-Cas9 half-life and found that, after two weeks upon functionalization and storage at 4°C, the AuNR.NTA-Cas9 lost its endonuclease activity (Figure 3D), implying protein degradation. Consequently, experiments were concluded within a timeframe of one to two weeks upon functionalization.

**Figure 3.**
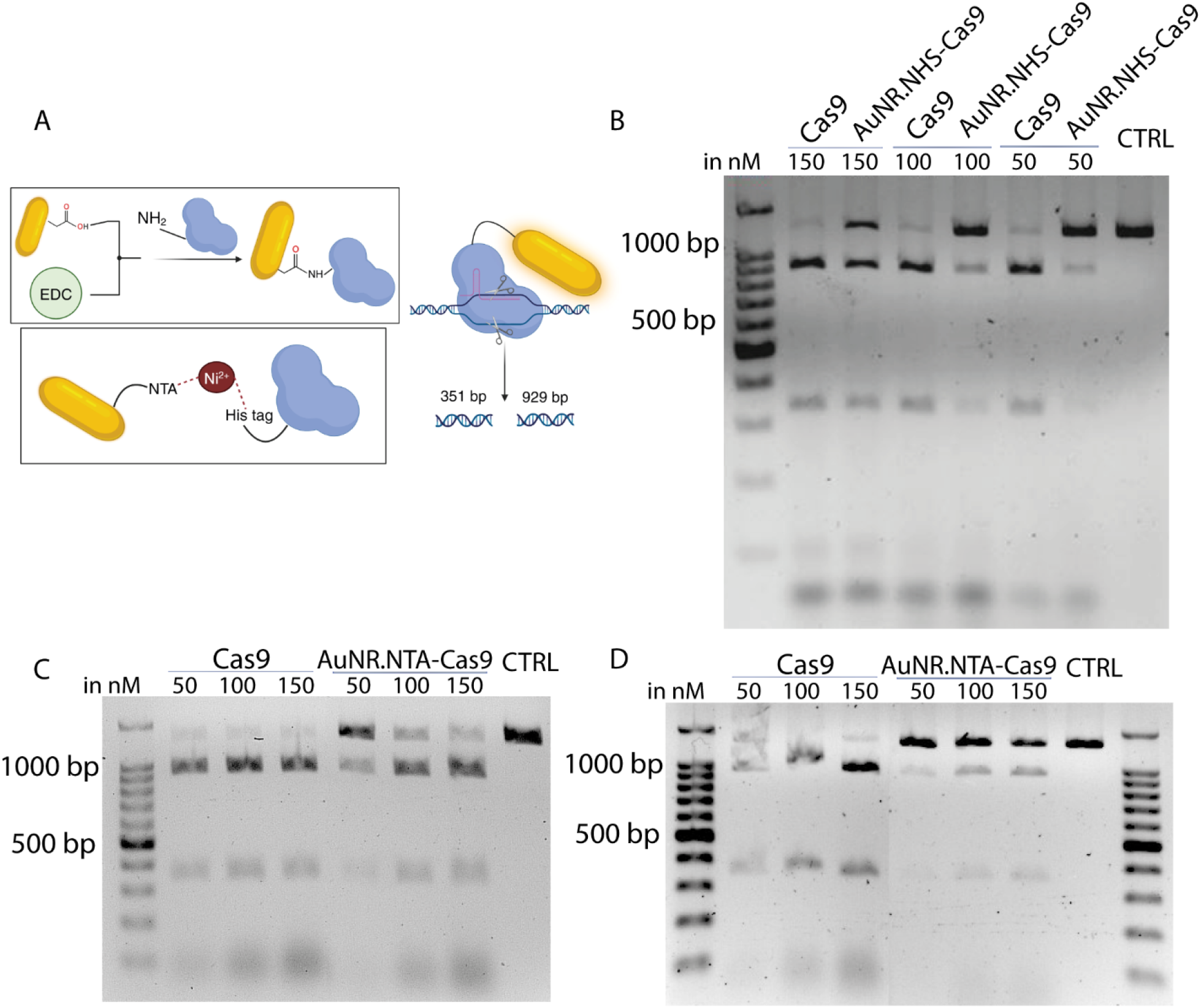
*In vitro* cleavage analysis of AuNR-Cas9 with different chemical backgrounds; NHS (AuNR.NHS-Cas9) or NTA (AuNR.NTA-Cas9). (A) Schematic illustration of the synthesis of the nanoformulation AuNR-Cas9 with the two different chemical backgrounds. (B) *In vitro* cleavage of the AuNR.NHS-Cas9 the first week upon functionalization. (C) *In vitro* cleavage efficiency of the AuNR.NTA-Cas9 the first week upon functionalization, and (D) two weeks upon functionalization after storage at 4°C. (B-D) As positive control, the *in vitro* cleavage efficiency of unconjugated Cas9 was also tested.

Next, we assessed the gene editing capability of the AuNR-Cas9 in zebrafish, by injecting the nanoformulation in the zygote targeting the tyrosinase gene (TYR) and checking the larvae after 3 days (Figure 4A). The AuNR-Cas9 in complex with the gRNA induced embryo depigmentation (Figure 4B, C), with most larvae exhibiting mild depigmentation at 3 days post-fertilization (dpf), whereas injection with the Cas9:gRNA resulted in strong and mild depigmentation phenotypes. Gene editing was confirmed by High-Resolution Melting (HRM) analysis (Figure 4D). Non-functionalized nanorods (AuNR) and AuNR-Cas9 without gRNA failed to induce gene editing in zebrafish, as their melt curves were similar to the control group (Figure 4E). Furthermore, the DNA of 6 AuNR-Cas9-injected zebrafish larvae was analyzed by Sanger sequencing and ICE software analysis, showing a mutagenesis efficiency ranging between 28-95% (Figure 4F), and inducing as predominant indel mutation a deletion of 4 nucleotides, D4 (GAG----GATA) (Figure 4G).

**Figure 4.**
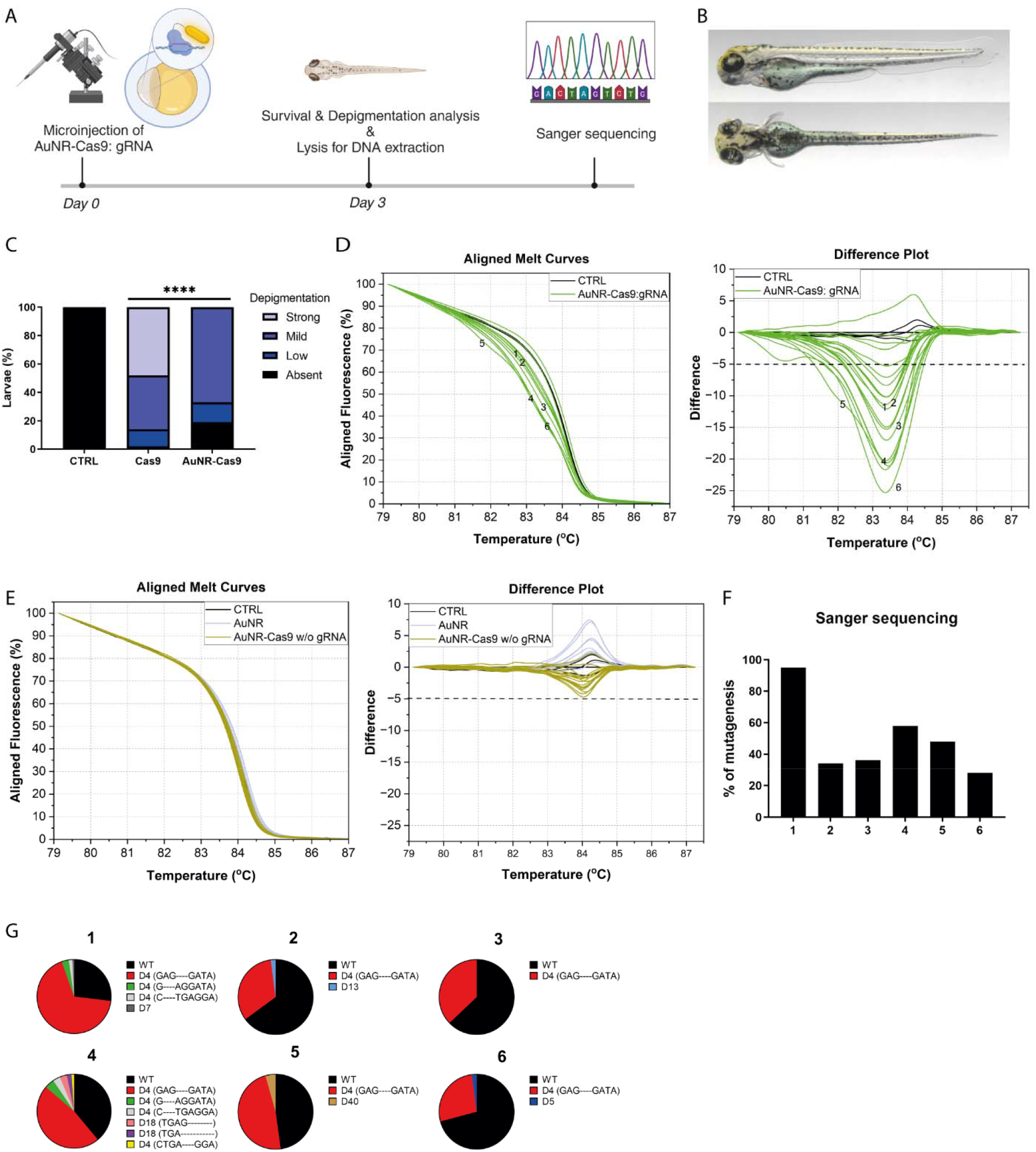
Gene editing efficiency of the AuNR-Cas9 in zebrafish. (A) Schematic illustration of the experimental process. (B) Representative images of depigmented zebrafish larvae injected with AuNR-Cas9: gRNA. (C) Quantification of depigmented phenotypes in zebrafish larvae injected with 750pg of Cas9: gRNA or AuNR-Cas9: gRNA; Cas9: gRNA N=15; AuNR-Cas9:.gRNA N=111. A χ2 test was performed p<0.0001. The microinjections of the zebrafish were performed in duplicate. (D - E) Melting analysis performed by High Resolution Melt Software (Applied Biosystems); for the AuNR-Cas9: gRNA N= 21, for AuNR N= 10, and AuNR-Cas9 without gRNA N= 10. (F - G) Sanger sequencing analysis by ICE software from Synthego. (F) Percentage of mutagenesis in each tested sample. (G) Analysis of the obtained indel mutations for each tested sample.

After demonstrating that the catalytic activity of Cas9 is not impaired by the functionalization, we conjugated the dCas9 protein onto gold nanorods following the pre-selected method, via affinity binding. The mechanism of action of our nano-switch requires the dimerization of two gold nanorods of AuNR-dCas9:gRNA (Figure 5A). To select the pair of gRNA at desired distances and orientation, we designed the I-GENEMatcher software that allows for selections of pair of gRNAs in target genes in 3 available genomes (Homo sapiens, Danio rerio, Mus musculus) upon analysis by commonly used software for the design of gRNAs (CHOPCHOP and CRISPRscan) (https://i-gene.d4science.org/group/i-genepublic/i-gene-tool). Using the I-GENEMatcher, we selected a pair of gRNAs targeting the first exon of the TYR gene in zebrafish, Danio rerio (danRer10 or danRer11), consisting of gRNA1 and gRNA2 in a distance of 9 nt as indicated by the I-GENEMatcher software in a PAM OUT configuration (Figure 5A). The AuNR-dCas9:gRNA1 and AuNR-dCas9:gRNA2 were incubated with a DNA linear fragment containing the first exon of the TYR gene. The sample was dried and analyzed by transmission electron microscopy (TEM). Figure 5B reveals the formation of dimers of AuNR onto a linear DNA fragment driven through the respective dCas9:gRNA1 and dCas9:gRNA2 ribonucleoproteins.

**Figure 5.**
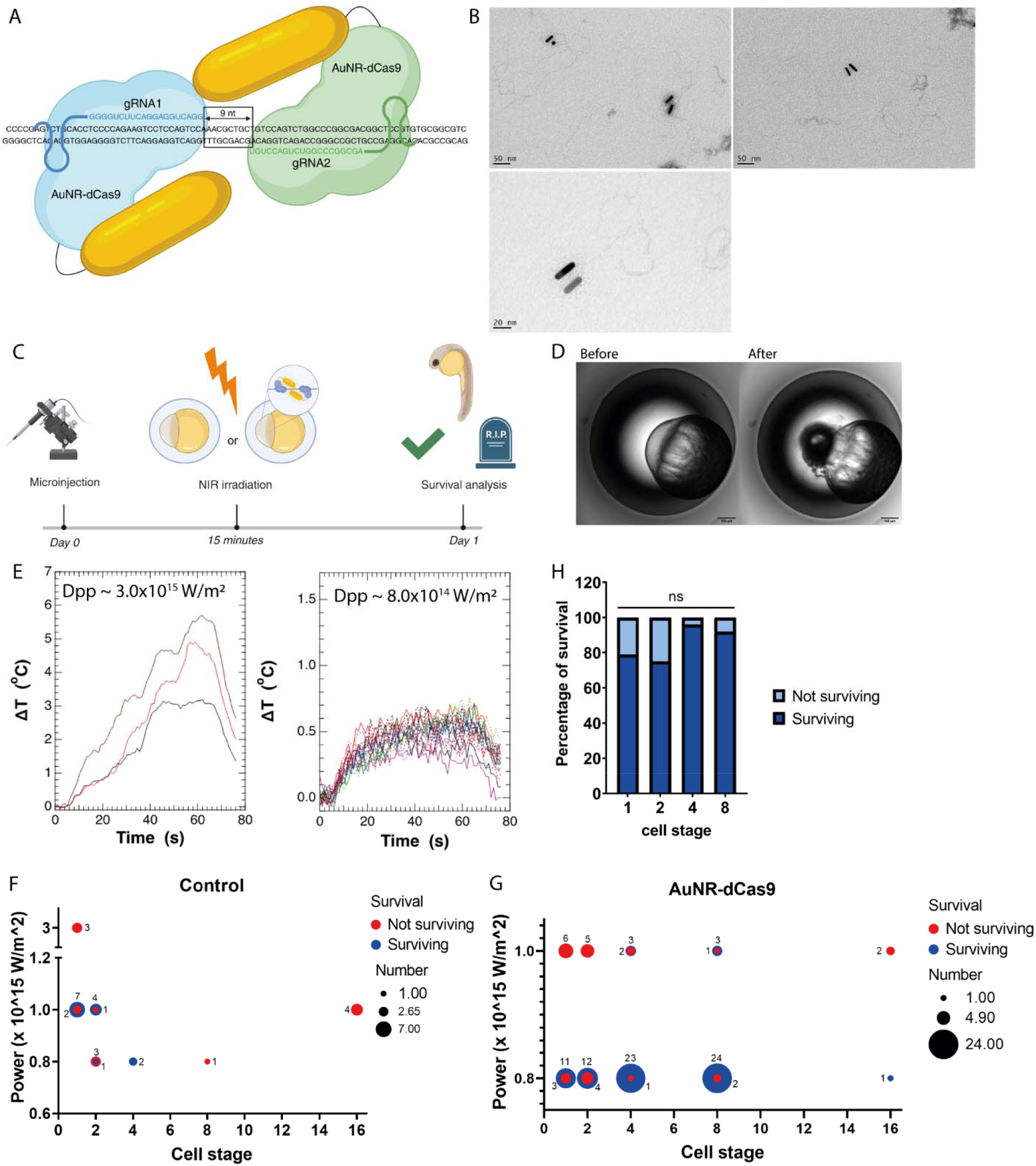
Testing the irradiation scheme of the dimer *in vivo*. (A) Schematic illustration showing the binding of two nanotransducers in proximity in the target region. (B) TEM micrographs showing the formation of dimers of gold nanorods onto a linear DNA fragment driven through the respective monomers dCas9:gRNA1 and dCas9: gRNA2. (C) Schematic illustration of the experimental work-flow. (D) Control uninjected zebrafish embryos at the 1-cell developmental stage showing laser ablation after irradiation at 3×10^15^ W/m^2^. (E) ΔT of the bulk temperature of embryos during irradiation. Embryos irradiated at 3×10^15^ W/m^2^ at the 1-cell stage are presented in the left panel, and at 8×10^14^ W/m^2^ in different cell stages in the right one. (F) Survival profile of control uninjected zebrafish embryos 1 day post-irradiation. (G) Survival profile 1-day post-irradiation of zebrafish embryos injected with the AuNR-dCas9: gRNA tyr1 and tyr5. (H) Survival percentage 1-day post-irradiation of zebrafish embryos injected with the AuNR-dCas9:gRNA1/gRNA2 and irradiated at 8×10^14^ W/m^2^ at different cell stages. Statistical analysis was performed by Fisher’s exact test; comparing 1- and 2-cell stage p>0.9999, comparing 1- and 4-cell stage p=0.1319, comparing 1- and 8-cell stage p=0.3223

Based on the peak energy density values suggested by simulations, we developed a specific experimental strategy aimed at improving the effectiveness of in-vivo irradiation experiments. To this aim, we decided to implement a spiral motion of the sample stage in order to reduce the effects of error-prone repair by extending the activation-irradiation time and to prevent the need for an exact co-localization of the DNA-nano-switch site with the focal spot of the laser. Moreover, we increased the laser power to bring above the activation threshold a sample volume of the order of 1 nl, while remaining below the limit for photothermal bubble generation ^11, 12^ or photo-toxicity effects. In order to assess the light activation of the nano-switch *in vivo* using zebrafish embryos, we first optimized the irradiation set-up. Zebrafish embryos control (uninjected) or injected with the AuNR-dCas9:gRNA1/gRNA2 in the zygote were exposed to a 200 fs laser at 870 nm and the survival of the irradiated zebrafish was evaluated 1 day post-fertilization (Figure 5C). To determine the optimal laser irradiation protocol, increasing peak power densities 8×10^1^□, 1×10^1^□, and 3×10^1^□ W/m^2^ were tested and delivered in a focal spot with waist of 5 µm, all above the value suggested by the simulations in order to compensate for actual local optical properties of the sample, e.g., nanorod alignment with respect to laser polarization, diffusion and scattering effects of the material. At the highest power density, the embryos were adversely affected, with cells likely destroyed due to photothermal bubble generation (Figure 5D). The bulk temperature increase (ΔT) during irradiation reached up to 5°C (Figure 5E), and none of the embryos irradiated at 3×10^1^□ W/m^2^ survived until 1-day post-irradiation (1 dpi) (Figure 5F). Consequently, we tested the lower power densities on zebrafish treated with AuNR-dCas9:gRNA1/gRNA2, to assess irradiation-induced toxicity. Embryos irradiated at 1×10^1^□ W/m^2^ also did not survive until 1 dpi (Figure 5G). Therefore, we selected the lowest power density, 8×10^1^□W/m^2^, which resulted in negligible ΔT (Figure 5E) and high survival rates (Figure 5G). Specifically, irradiating at 8×10^1^□ W/m^2^ at the 1-cell stage yielded a higher than 75% survival rate (Figure 5H). Though, survival rates improved when irradiation was performed at later developmental stages, such as the 4- and 8-cell stage, there was no statistical difference in survivability compared to the 1-cell stage (Figure 5H).

As we mentioned earlier, we employed two different irradiation schemes, referred to as “3D-spiral” and “2D-spiral” (Figure 6A). In both methods, the laser power density at the focal spot was maintained at 8x101^4^ W/m^2^, with the spot moving in a spiral pattern to cover an area with a 150 μm radius over 60 seconds. The primary distinction between the two protocols is the movement of the focal plane in the “3D-spiral” scheme, which affects the distribution of laser energy within the irradiated volume. In the “2D-spiral” scheme, the focal plane remains fixed at a set distance below the surface of the sample, typically around 50 μm, which approximately corresponds to the Rayleigh range under the experimental conditions. Conversely, in the “3D-spiral” scheme, the focal plane shifts within the cell volume during irradiation. The total irradiation time of 60 seconds is divided into four equal phases, with the laser spot moving along a spiral path in each phase. After completing a spiral, the focal plane shifts by 10 μm, and the spiral movement repeats (Figure 6A).

**Figure 6.**
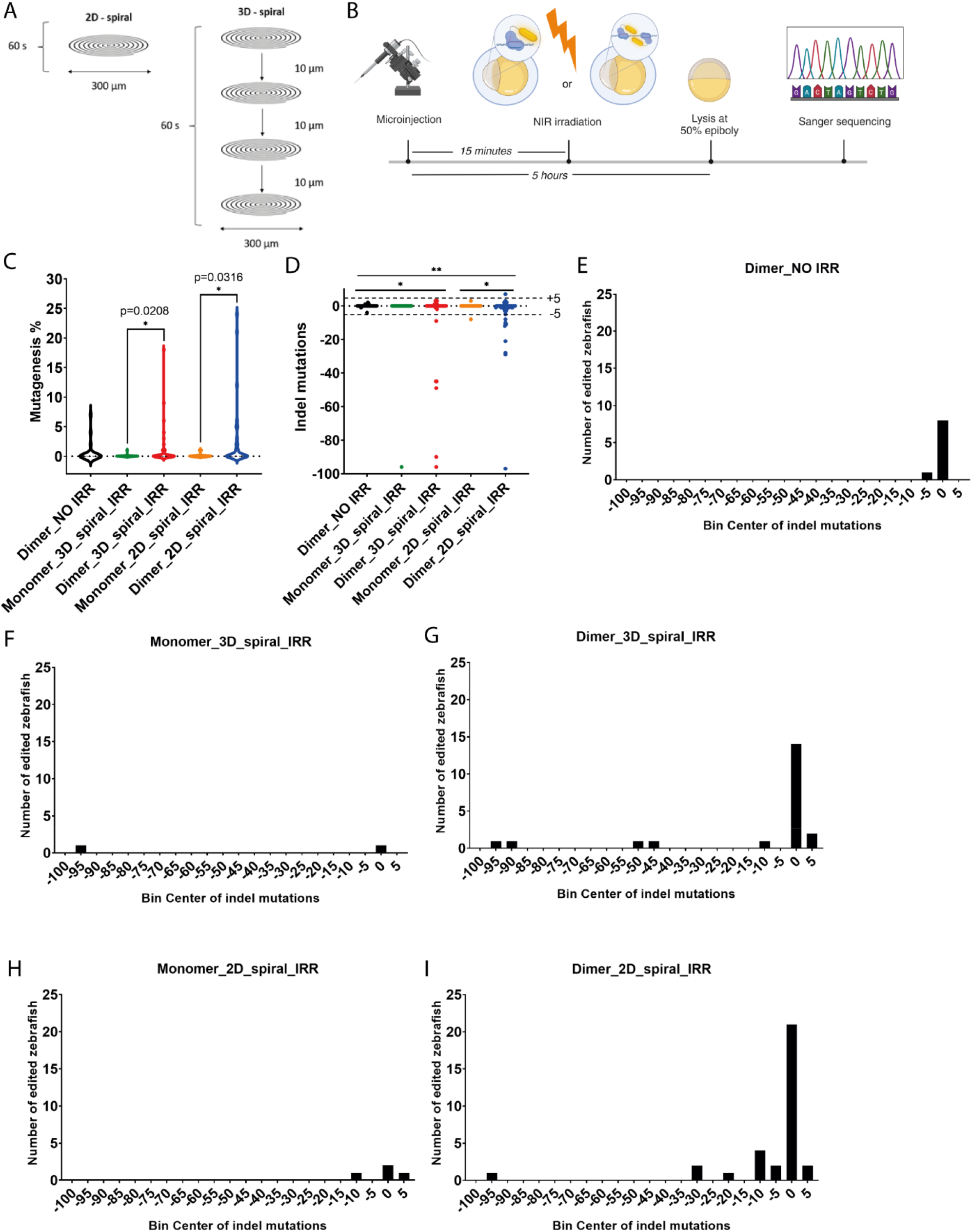
Gene editing analysis on the tyrosinase target gene in zebrafish upon on-demand light-activation of the nano-switch. (A) Schematic illustration of the two irradiation schemes; “2D-spiral”, and “3D-spiral”, showing the movement of the laser focal spot in 60 seconds of irradiation. (B) Schematic illustration of the experimental workflow. (C) Mutagenesis percentages of the tested zebrafish embryos injected with AuNR-dCas9:gRNA1/gRNA2 (Dimer) or only AuNR-dCas9:gRNA1 (Monomer) and then irradiated (IRR) with 2D-spiral (2D) or 3D-spiral (3D) scheme. As control, zebrafish zygotes were only injected with AuNR-dCas9:gRNA1/gRNA2 but not irradiated (Dimer_NO IRR). The analysis was performed by the Mann-Whitney test. The ns difference is not represented in the plot. N for Dimer_NO IRR was 24, N for Monomer_3D_spiral_IRR was 24, N for Dimer_3D_spiral_IRR was 32, N for Monomer_2D_spiral_IRR was 20, N for Dimer_2D_spiral_IRR was 36. The analysed groups were Dimer_NO IRR vs Dimer_3D_spiral_IRR (p=0.1937) and Dimer_2D_spiral_IRR (p=0.0846), Monomer_3D_spiral_IRR vs Dimer_3D_spiral_IRR (p=0.0208), and Monomer_2D_spiral_IRR vs Dimer_2D_spiral_IRR (p=0.0316). (D) Indel mutations of the tested zebrafish. Data were analyzed by Chi-square test. The ns difference is not represented in the plot. N for Dimer_NO IRR was 28, N for Monomer_3D_spiral_IRR was 24, N for Dimer_3D_spiral_IRR was 37, N for Monomer_2D_spiral_IRR was 20, N for Dimer_2D_spiral_IRR was 52. The analysed groups were Dimer_NO IRR vs Dimer_3D_spiral_IRR (p=0.0327) and Dimer_2D_spiral_IRR (p=0.0016), Monomer_3D_spiral_IRR vs Dimer_3D_spiral_IRR (p=0.1492), and Monomer_2D_spiral_IRR vs Dimer_2D_spiral_IRR (p=0.0293). (E-I) Plotted histograms of the N of indel mutations of every reported group.

To assess the gene editing efficiency of the nanotransducer AuNR-dCas9 on the target DNA locus, we injected the AuNR-dCas9: gRNA1/gRNA2 in the zebrafish zygote. After approximately 15 minutes, the injected zebrafish underwent the irradiation process with either of the two previously mentioned schemes, “3D-spiral” or “2D-spiral”, at different developmental stages (1-, 2-, 4- or 8-cell stage). After the irradiation, the zebrafish were placed at 29°C and were lysed at 5 hours post fertilization (hpf), at the stage of 50% epiboly (Figure 6B). The hypothesis is that upon irradiation at 870 nm, the dimer of the gold nanorods is activated leading to extremely high localized heat that would be subsequently responsible for a thermo-inducible DSB. Then, the break has to be repaired, resulting in indel mutations induced by error-prone DNA repair mechanisms.

Zebrafish zygotes were injected with the AuNR-dCas9:gRNA1/gRNA2, to create the dimer formation, and were irradiated with either the scheme of “3D-spiral” or “2D-spiral” which both led to indel mutations and/or fragment deletions in the target DNA. In detail, the DNA of zebrafish irradiated with the “3D-spiral” showed 28.13% mutagenesis including mostly mutations corresponding to fragment deletions, in the bin center of -45 to -50 and -90 to -95 (Figure 6G). In this case, the mutagenesis efficiency ranged between 1-18% per sample (Figure 6C). On the other hand, when the zebrafish were irradiated with the scheme “2D-spiral”, they showed 33.33% mutagenesis including a high variety of indel mutations per sample (Figure 6I) and a higher efficiency of mutagenesis per sample than with the scheme “3D-spiral” reaching up to 24% (Figure 6C). All the sequencing results are provided as Supporting Information.

Zebrafish that were injected with the monomer and irradiated at 870 nm sporadically presented indel mutations (Figure 6F, H). This result is not surprising because even the monomer shows some residual optical absorption at the used wavelength, and it can be rarely activated with a very low percentage of mutagenesis. Indeed, comparing the percentage of mutagenesis between the monomer and the dimer for both irradiation schemes, there is a statistical difference of p=0.0208 for the “3D-spiral” irradiation scheme and of p=0.0316 for the “2D-spiral” irradiation scheme (Figure 6C). Furthermore, it was shown that there is a significant presence of indel mutations in zebrafish injected with the dimer and irradiated with either the “3D-spiral” (p=0.0327) or the “2D-spiral” (p=0.0016) compared to the zebrafish that were injected with the AuNR-dCas9: gRNA1/gRNA2 but not irradiated (Figure 6D) despite the presence of small indel mutations in the latter group (Figure 6E

## CONCLUSION

In this paper, we provide a proof-of-concept of a new technology for spatiotemporal control of inducible<u>-gen</u>ome <u>e</u>diting (I-GENE technology). Specifically, we generated a nano-switch that is composed of a sensor and a nanotransducer. The sensor is the DNA recognition element, i.e., the dCas9:gRNA that is responsible for guiding the nano-switch in the desired genomic location. The nanotransducer is a plasmonic nanoparticle that is highly tuned to absorb a specific optical wavelength and efficiently convert it into heat. In this system, the actuator is the heat generated by the nano-switch under irradiation that is used for a thermo-inducible DNA DSB. Numerical analysis pointed out that heat generation can be more efficient when the metallic core is in the form of nanorods. Moreover, gold nanorods (AuNRs) of 5 nm diameter and 20 nm length have an optical absorption peak of 780-810 nm in the form of monomer and 850-910 nm in the form of dimer. Indeed, irradiating at the wavelength of 870 nm, the dimer (but not the monomer) is activated, leading to an extreme increase in temperature (>150 °C) at the zeptoliter volume surrounding the dimer, sufficient for determining a thermo-inducible DSB. The monomer was generated by conjugated Cas proteins to the AuNR His-tag affinity. This functionalization scheme preserves the biological activity of the protein. Using a pair of guides having their DNA target sequence respectively on each filament, spaced 9 nt apart and in a PAM OUT configuration, we validated the ability of the nano-switch to localize as a dimer on the target DNA locus guided by the pair of gRNA1/gRNA2. The nano-switch was validated in zebrafish embryos injected with the pair AuNR-dCas9:gRNA1/gRNA2 and irradiated with a wavelength of 870 nm in pulsed mode (200 fs pulses at 80 MHz repetition rate) and local average power density of 2×10^9^ W/m^2^ We found that the dimer induces a statistically significant increase in the indel mutation in comparison to the irradiated monomer and the non-irradiated dimer.In conclusion, inspired by the principle of synthetic biology, we generated a nano-switch that implements a multi-input AND logic gate that produces a single binary output (gene editing) conditioned by a number of binary inputs (a resonant light, loci recognition of the gRNAs, dimerization).

## IMITATION OF THE STUDY

A limitation of this study was the inability to measure the local temperature increase in a molecular level. We attempted to synthesize an intracellular temperature sensor^13^ but we were unsuccessful in obtaining reproducible measurements of the temperature increase within the zeptoliter volume of the particles.

## MATERIAS AND METHODS

### Data and metadata are available in ZENODO (10.5281/zenodo.11640533). Simulations

Commercially available COMSOL Multiphysics® software was used to evaluate the optical properties of the AuNPs (absorption, scattering, local electric field intensity, and electromagnetic power loss density) and estimate their local heating exploited for DNA strand break. The refractive index and extinction coefficient of gold were taken from the work of Rakić et al.^14^,and the refractive index of the environment (water) was set to 1.33.

### Nanotransducer synthesis

Gold nanorods were bought from Nanopartz, C12-5-780-TC-DRY-2.5 and C12-5-780-TNT-PBS-50-1. The dCas9 protein was bought from IDT, Alt-R® S.p. dCas9 protein V3, # 1081067. For the NHS chemistry protocol, a solution containing 2×10^13^ nanoparticles (C12-5-780-TC-DRY-2.5) is incubated for 20 minutes at room temperature with 1-ethyl-3-(3-dimethylaminopropyl) carbodiimide hydrochloride (EDC) (PG82073, Thermo Scientific) and N-hydroxysulfosuccinimide (Sulfo-NHS) (PG82071, Thermo Scientific) in 1 ml of 2-(N-morpholino)ethanesulfonic acid (MES) buffer at pH 6. The mixture is transferred into a VIVAspin 500 centrifugal column (30000 MW, VS0122, Sartorius) and washed twice in PBS. The resulting solution is then concentrated to 500 μl and placed in a glass vial to incubate with 50 μg of protein for one hour at room temperature. After protein conjugation, 50 mM Tris at pH 8.8 is added for 10 minutes. The functionalized nanoparticles are subsequently washed three times by centrifugation, resuspended in 40 μl of PBS, and stored at 4°C. For the NTA chemistry protocol, a solution containing 2×10^13^ nanoparticles (C12-5-780-CUS-PBS-50-1) is incubated with 50 μg of protein in PBS for one hour at room temperature in a glass vial. Following protein conjugation, the functionalized nanoparticles undergo three washes by centrifugation, are resuspended in 40 μl of PBS, and then stored at 4°C.

### In vitro cleavage assay

The DNA linear fragment included the entire first exon of the tyrosinase gene of zebrafish (Danio rerio) and was synthetized as part of a plasmid ordered by IDT. Using the CHOPCHOP software, the gRNA with sequence GGGCCGCAGTATCCTCACTC (5’-3’) was identified and used for the in vitro studies. The protocol was provided by IDT https://sfvideo.blob.core.windows.net/sitefinity/docs/default-source/protocol/alt-r-crispr-cas9-protocol-in-vitro-cleavage-of-target-dna-with-rnp-complex.pdf?sfvrsn=88c43107_30. The cleavage sample was combined with a 6X loading buffer (R0611, Thermo Scientific) and loaded onto a 1.5% agarose gel. A 100 bp DNA ladder (G210A, Promega) served as a reference. The signals were visualized using the ChemiDoc™ XRS+ Imaging system (BioRad), and the images were obtained using ImageLab 6.1 software.

### Zebrafish

All animal procedures were conducted in strict accordance with protocols approved by the Italian Ministry of Public Health and the local Ethical Committee of the University of Pisa (authorization no. 99/2012-A, dated 19.04.2012), in compliance with Directive 2010/63/EU. Zebrafish were bred in the animal facility of the University of Pisa (Authorization Number DN-16 /43 on 19/01/2015, renewal Authorization Number 1695 on 12/10/2023N).

Zebrafish were microinjected according to the protocol https://sfvideo.blob.core.windows.net/sitefinity/docs/default-source/user-submitted-method/crispr-cas9-rnp-delivery-zebrafish-embryos-j-essnerc46b5a1532796e2eaa53ff00001c1b3c.pdf?sfvrsn=52123407_4,using the Pneumatic Picopump PV820 air microinjector (World Precision Instruments). For the AuNR-Cas9 experiments, the same gRNA as for the in vitro studies was used. The pair of gRNAs to form the dimerization of the nanotransducers was selected using the I-GENEMatcher software, gRNA1: TGTCCAGTCTGGCCCGGCGA, and gRNA2: CCCCAGAAGTCCTCCAGTCC.

The zebrafish embryos were kept in E3 water (5 mM NaCl, 0.17 mM KCl, 0.33 mM CaCl2, 0.33 mM MgSO4) at 29°C. The development was controlled till 3 days post-fertilization (dpf). Zebrafish larvae were anesthetized with tricaine 1X (A5040, SIGMA-ALDRICH) for acquiring images at 3 dpf using the Nikon stereomicroscope SMZ1500 n. At 3dpf, zebrafish were lysed in 20 μl Thermopol buffer (B90004S, New England Biolabs) and heated at 95°C for 10 minutes. Then, 5 μl of Proteinase K solution (20mg/ml) was added to each sample, incubating them at 55°C for 1 hour and then at 95°C for 10 minutes to inactivate the enzymatic activity.

For the High-Resolution Melting, the following protocol was used https://assets.thermofisher.com/TFS-Assets/LSG/manuals/cms_069853.pdf. In the Difference plots, a threshold was indicated at -5 considering the profile of the control groups. Significant lower melting profiles were considered presenting a difference <-5.

Furthermore, a standard PCR protocol was used to amplify the genomic DNA and the amplicon was purified by Monarch® PCR & DNA Cleanup Kit from New England BioLabs Inc (#T1030S). The Sanger sequencing was performed by Genewiz. Analysis of the sequencing data was performed with the ICE software by Synthego. To define the presence of indel mutations by analyzing the ICE results, indel mutations in the range of -5< x <+5 were considered as “false-positive” mutations and were excluded from the category of indel mutations.Indel mutations >+5 and <-5, and Fragment deletions (as indicated from the analysis with the ICE software) were considered in a common category.

**Table 1.**
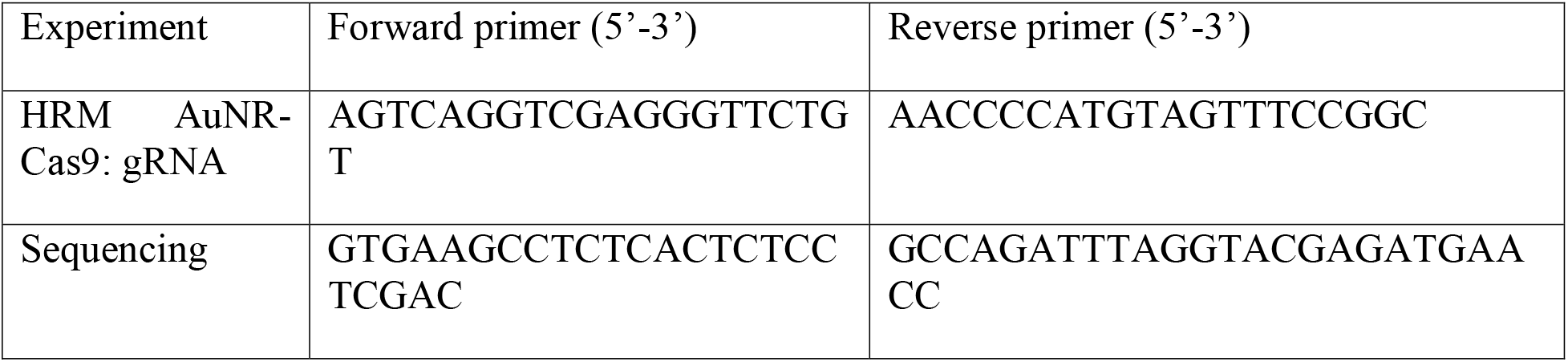
List with primers.

### Transmission Electron Microscopy

The formation of dimers of gold nanorods onto a linear DNA fragment morphology was analyzed through transmission electron microscopy (TEM), with a Jeol JEM 1011 electron microscope using an acceleration voltage of 100 kV and equipped with a 2 Mp charge-coupled device (CCD) camera (Gatan Orius SC100). A drop of the samples was deposited on a copper grid of 150 mesh, previously coated with an amorphous carbon film and plasma treated to remove hydrocarbon residues. The samples were then stained, treating the grids with a 1% uranyl acetate solution in water for 30 s before the specimen dried.

### Irradiation

The used laser source was a tunable ultrafast Ti:Sapphire laser, model Sprite-XT from the M2 company. Power was adjusted by a rotatable waveplate/polarizer assembly and measured by a Scientech 361 power meter with a 10% accuracy. The laser beam was expanded by a 1:2 telescope, coupled to a Nikon Eclipse Ti2-E microscope by the rear port and focused onto the sample with a 45-degree mirror and a Nikon CFI E-Plan 10x, 0.25 NA objective

One zebrafish embryo was deposited as a drop in E3 water on a glass coverslip which was located on a Nikon Ti2-S-SS-E motorized scanning stage controlled by NIS Element AR software and a National Instruments NI-PCIe-6321 card. An infrared temperature detector (Melexis MLX90614xCI) was positioned above the drop to monitor and record the bulk temperature of the entire sample during irradiation. Irradiation begins at the center of the sample, which is automatically identified, and the stage moves in a spiral pattern.

## ASSOCIATED CONTENT

## Supporting Information

The Supporting Information is available free of charge.

Supporting video: Temperature profile at the surface of the two nanorods (avi)

Supporting figures: Sanger sequencing analysis results by the ICE software for every sequenced sample (PDF)

## AUTHOR INFORMATION

## Author Contributions

Conceptualization: V.R., F.T, F.F.; methodology: V.R., F.T, F.F., C.G, S.K., software: F.T.,F.D.A., S.K.; validation: S.K., M.A.R., F.N., D.D., M.D.M., F.F.; formal analysis: S.K., V.R.; Data curation: S.K., V.R.; Writing - Original Draft: S.K., V.R.; Writing - Review & Editing: All; Supervision: V.R., F.T., F.F., C.G.; Funding acquisition: V.R., F.T., F.D.A.

## ACKNOWLEDGMENT

The authors warmly acknowledge Aliaksandr Hubarevich for his important support in compiling the numerical simulations and his contribution to the article. The authors thank Nubisware for the technical support with the I-GENE Matcher software. The authors would like to thank Dr.

Elena Landi for her constant support.

Illustrations and experimental schemes were created with Biorender.com.

## ABBREVIATIONS

dCas9, dead Cas9; AuNR: gold nanorod
AuNR-dCas9: gold nanorod functionalized with the enzyme dCas9
LSPR: Localized Surface Plasmon Resonance
NHS: N-Hydroxysuccinimide
NTA: Nitrilotriacetic acid
HRM: High-Resolution Melting
TEM: Transmission Electron Microscopy

## TABLE OF CONTENTS GRAPHIC

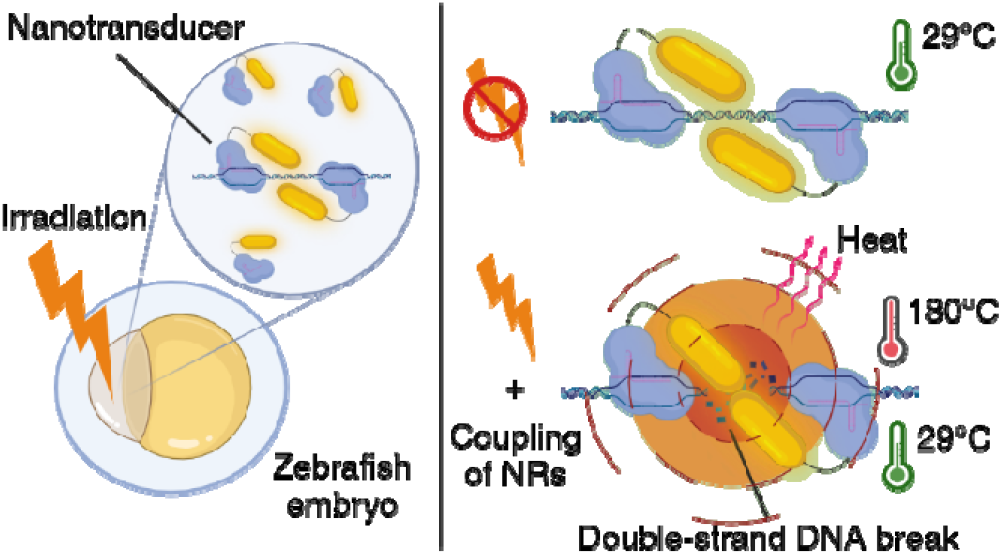

