## Supplementary data for "A photo-switchable gold nanoformulation based on the dCas9 protein for spatiotemporal controlled gene editing activation in vivo"

Supplementary figures

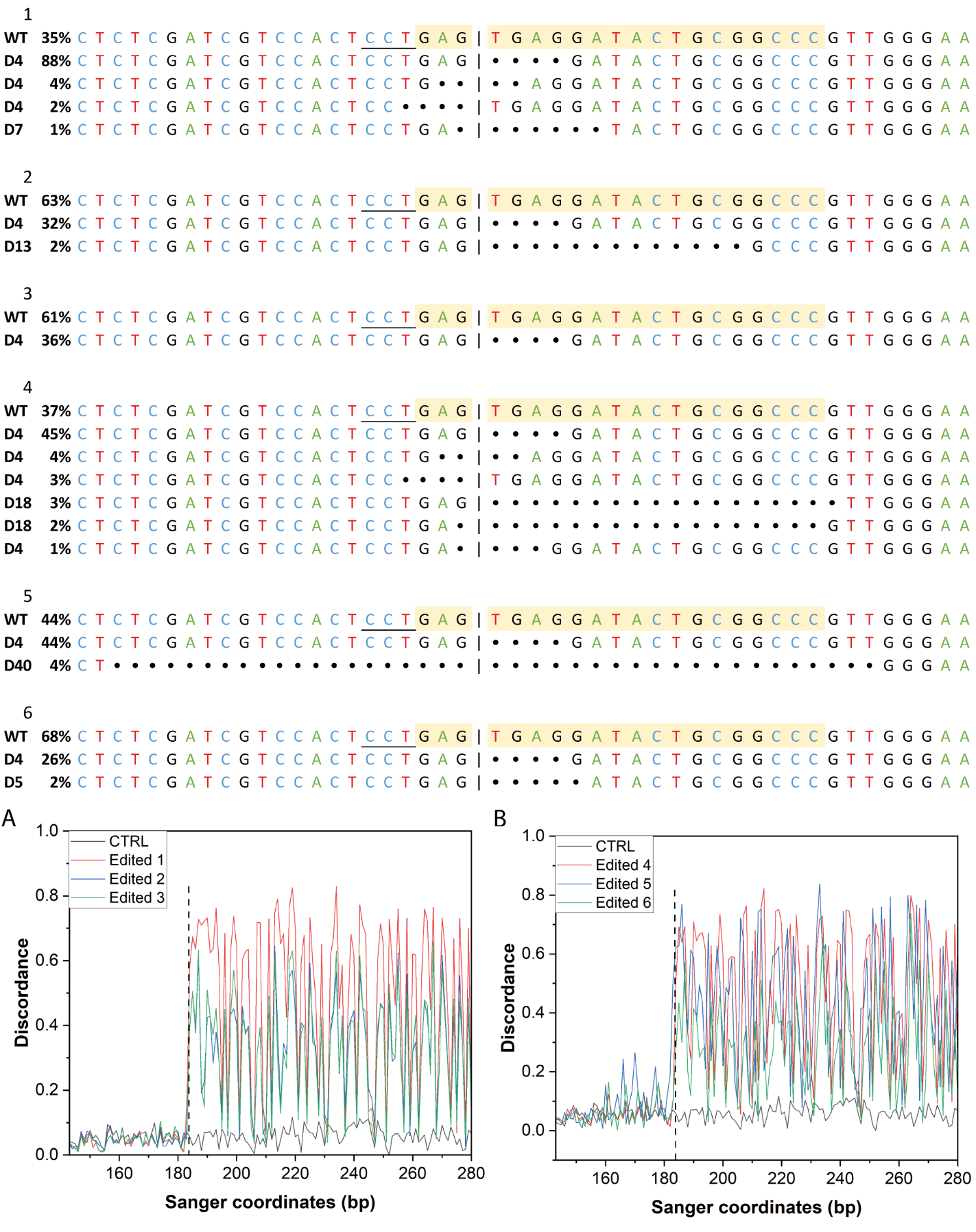

**Supplementary Figure 1**. Sanger sequencing analysis of zebrafish injected with the AuNR.NTA-Cas9: gRNA tyr2. Samples 1 to 3 correspond to the first experiment, and samples 4 to 6 to the second experiment. The targeted sequence by the gRNA tyr2 is highlighted in yellow. The line corresponds to the cutting site. The PAM sequence is underlined. Graphs A and B show the discordance of the edited samples compared to the control of the same experiment. The dotted line corresponds to the cutting site.

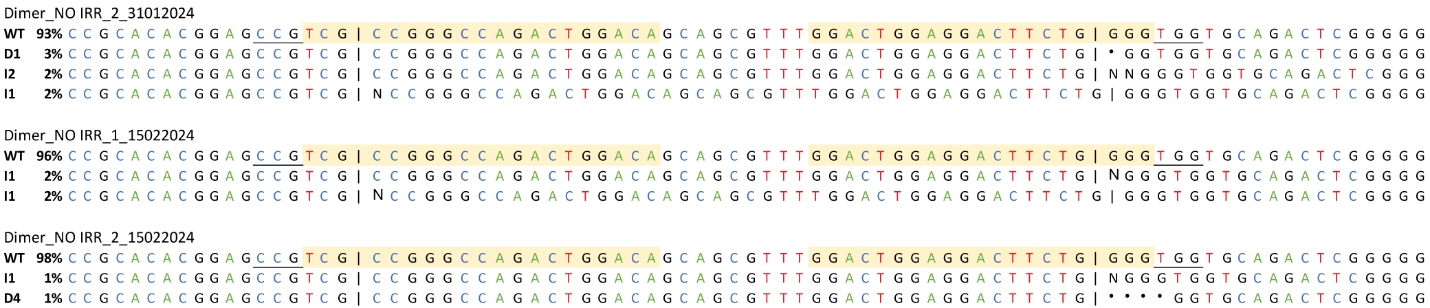

**Supplementary Figure 2.** Samples with indel mutations from zebrafish only injected with the AuNR-dCas9: gRNA tyr1 and tyr5, but not irradiatied (Dimer_NO IRR)

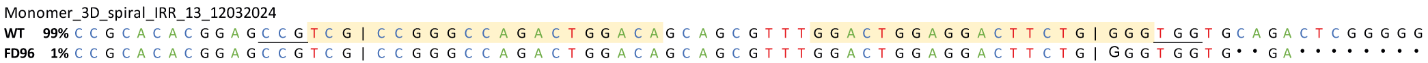

**Supplementary Figure 3.** Sample with indel mutations from zebrafish injected with the AuNR-dCas9: gRNA tyr5, and irradiatied with the scheme “3D-spiral” (Monomer_3D_spiral_IRR)

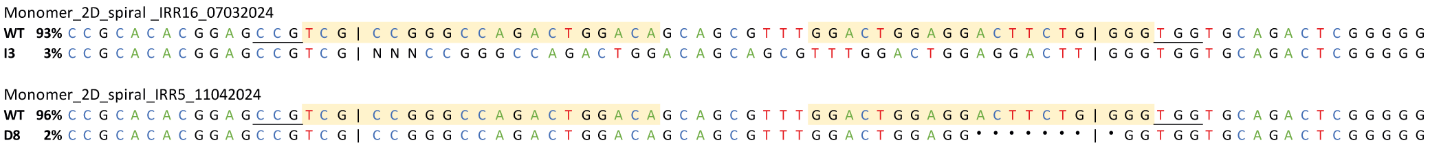

**Supplementary Figure 4.** Samples with indel mutations from zebrafish injected with the AuNR-dCas9: gRNA tyr5, and irradiatied with the scheme “2D-spiral” (Monomer_2D_spiral_IRR)

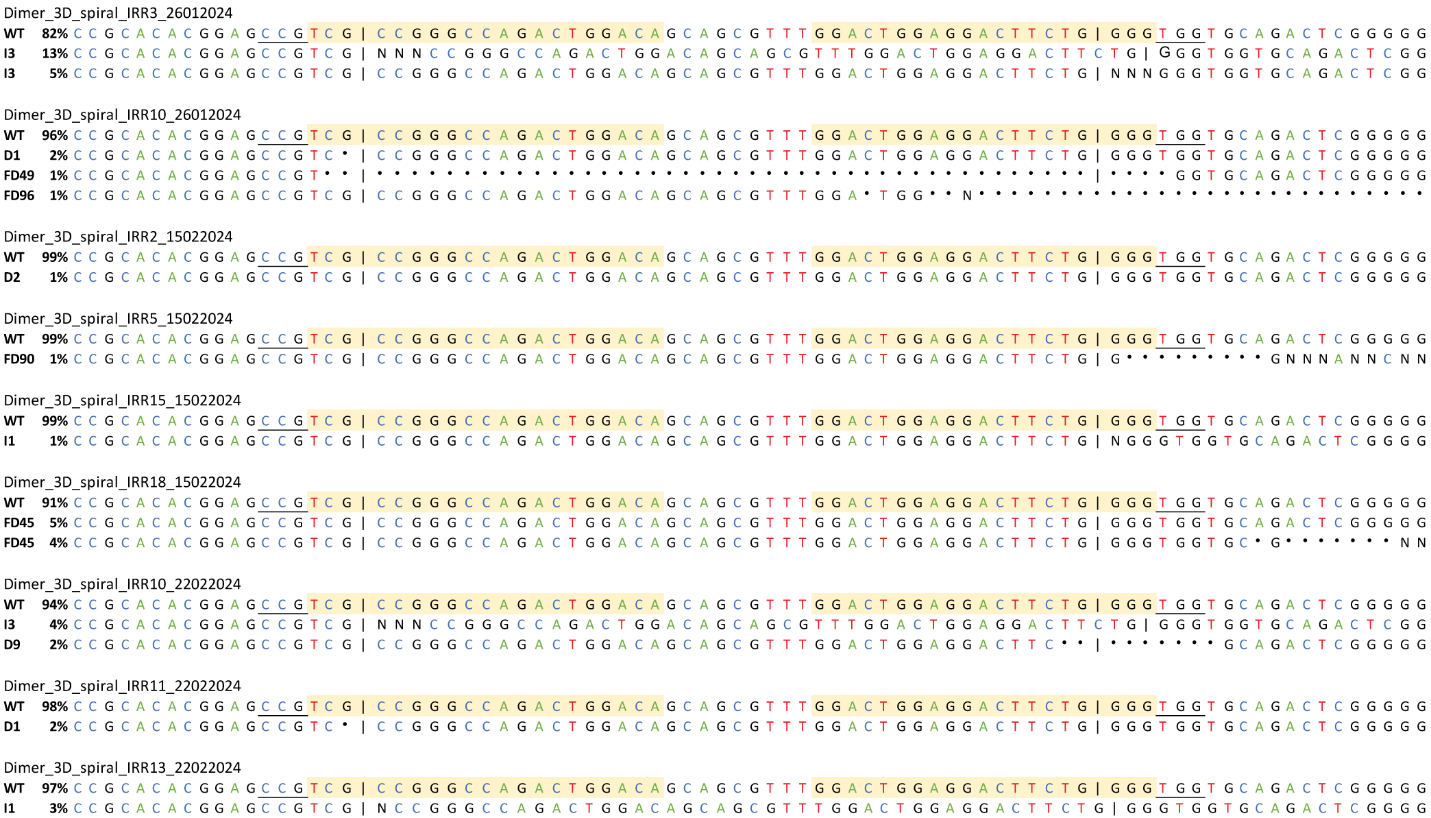

**Supplementary Figure 5.** Sample with indel mutations from zebrafish injected with the AuNR-dCas9: gRNA tyr1 and tyr5, and irradiatied with the scheme “3D-spiral” (Dimer_3D_spiral_IRR)

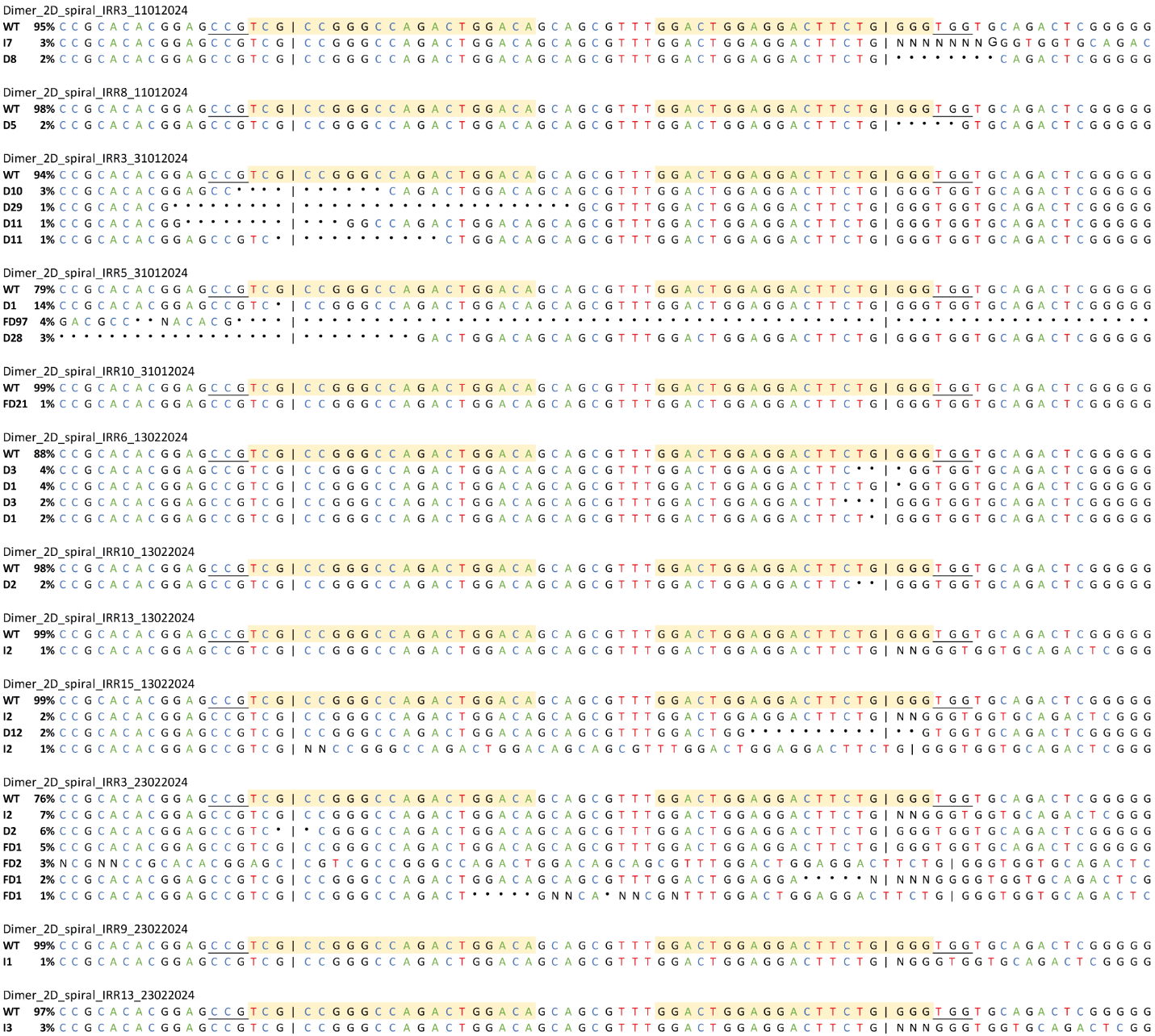

**Supplementary Figure 6.** Sample with indel mutations from zebrafish injected with the AuNR-dCas9: gRNA tyr1 and tyr5, and irradiatied with the scheme “2D-spiral” (Dimer_2D_spiral_IRR)
